# Cortical Acetylcholine Response to Deep Brain Stimulation of the Basal Forebrain

**DOI:** 10.1101/2024.07.30.605828

**Authors:** Khadijah Shanazz, Kun Xie, Tucker Oliver, Jamal Bogan, Fernando Vale, Jeremy Sword, Sergei A. Kirov, Alvin Terry, Philip O’Herron, David T Blake

**Author notes:** Correspondence to, Department of Neuroscience & Regenerative Medicine, 1120 15th Street, Rm. CA4002, Medical College of Georgia Augusta University, Augusta, GA 30912.

## Abstract

**Background:** Deep brain stimulation (DBS), the direct electrical stimulation of neuronal tissue in the basal forebrain to enhance release of the neurotransmitter acetylcholine, is under consideration as a method to improve executive function in patients with dementia. While some small studies indicate a positive response in the clinical setting, the relationship between DBS and acetylcholine pharmacokinetics is incompletely understood.

**Objective:** We examined the cortical acetylcholine response to different stimulation parameters of the basal forebrain.

**Methods:** 2-photon imaging was combined with deep brain stimulation. Stimulating electrodes were implanted in the subpallidal basal forebrain, and the ipsilateral somatosensory cortex was imaged. Acetylcholine activity was determined using the GRAB_ACh-3.0_ muscarinic acetylcholine receptor sensor, and blood vessels were imaged with Texas red.

**Results:** Experiments manipulating pulse train frequency demonstrated that integrated acetylcholine induced fluorescence was insensitive to frequency, and that peak levels were achieved with frequencies from 60 to 130 Hz. Altering pulse train length indicated that longer stimulation resulted in higher peaks and more activation with sublinear summation. The acetylcholinesterase inhibitor donepezil increased the peak response to 10s of stimulation at 60Hz, and the integrated response increased 57% with the 2 mg/kg dose, and 126% with the 4 mg/kg dose. Acetylcholine levels returned to baseline with a time constant of 14 to 18 seconds in all experiments.

**Conclusions:** These data demonstrate that acetylcholine receptor activation is insensitive to frequency between 60 and 130 Hz. High peak responses are achieved with up to 900 pulses. Donepezil increases total acetylcholine receptor activation associated with DBS but did not change temporal kinetics. The long time constants observed in the cerebral cortex add to the evidence supporting volume in addition to synaptic transmission.

**Graphical Abstract:** 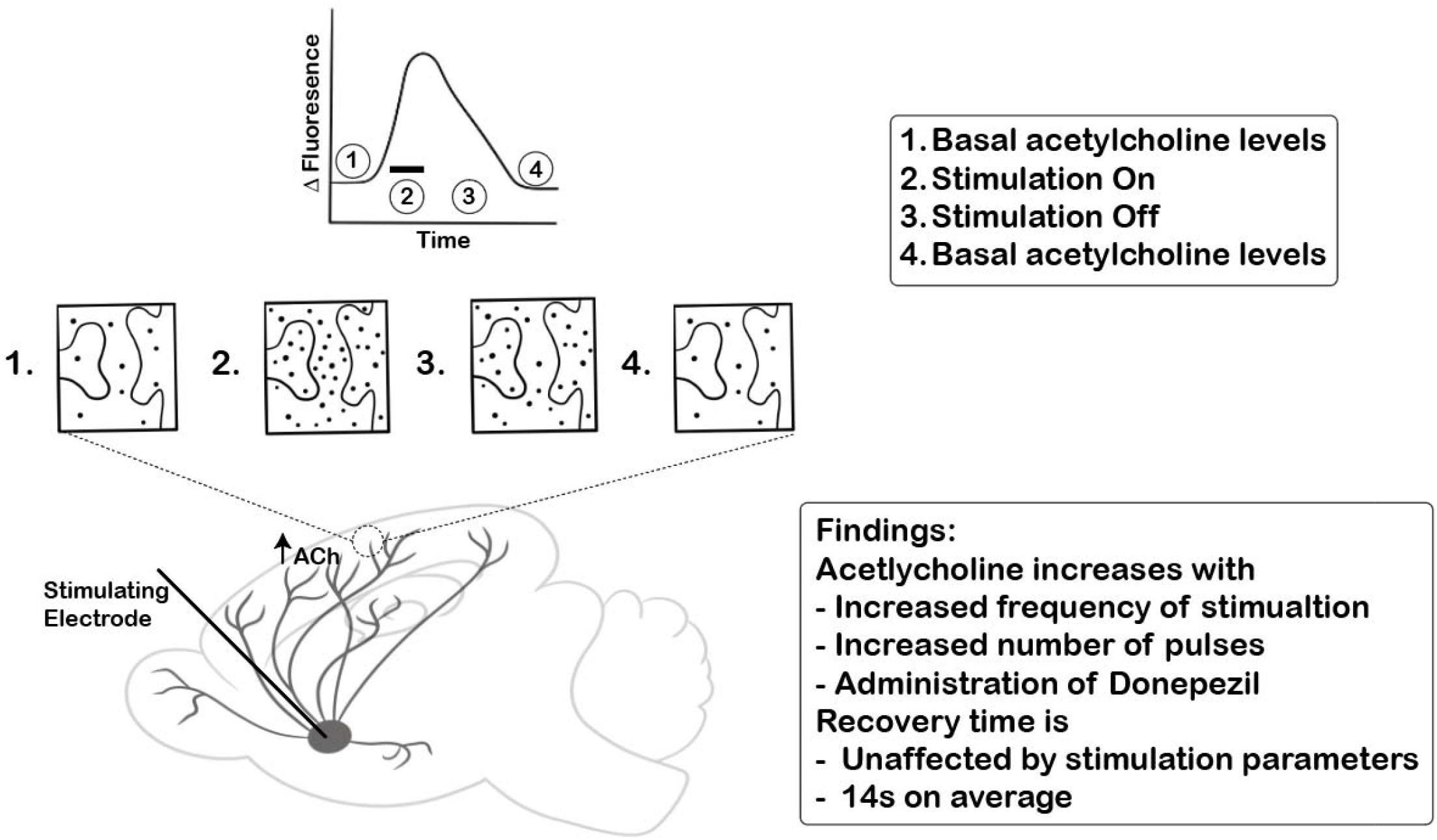

## Introduction

The basal forebrain is rich in cholinergic neurons that project axon terminals to the cortical mantle and release acetylcholine, a crucial neuromodulator implicated in cognitive processes and awake states [1–5]. Deficits in cholinergic neurotransmission in the brain are a hallmark of age-related cognitive decline and dementia [3,6–8] underscoring the importance of acetylcholine in maintaining cognitive function. For more than 30 years the standard treatment approach to attenuate these cholinergic deficits and improve cognitive function in dementia has been the use of acetylcholinesterase inhibitors (e.g., donepezil) which prevent the breakdown of acetylcholine [9]. This treatment approach provides modest efficacy, but is limited by dose dependent side effects that can be both centrally and peripherally mediated [10]. Importantly, electrical stimulation of the basal forebrain has been shown to improve working memory in primates [11] and spatial learning in rodent models of Alzheimer’s Disease [12]. This approach provides a method of selectively enhancing acetylcholine release in the brain, thus avoiding peripheral side effects. Theoretically, electrical stimulation of the basal forebrain might serve as a replacement for acetylcholinesterase inhibitors (AChEIs), or as an adjunctive treatment with AChEIs to further elevate acetylcholine levels since therapeutic doses of AChEIs typically result in only about 40% inhibition of the enzyme in the brain [13]. Despite clear efficacy of electrical stimulation of the basal forebrain in attenuating decrements in cognitive function and improving cholinergic tone, the pharmacokinetics of acetylcholine release in response to electrical stimulation remain poorly understood. A better understanding of the dynamics of acetylcholine release associated with different deep brain stimulation (DBS) parameters could facilitate both the DBS standalone treatment approach as well as the adjunctive treatment strategy described above (i.e., DBS + AChEI).

The acetylcholine response to basal forebrain stimulation has been studied for decades. Studies using deep brain stimulation and microdialysis in mice hippocampus show peak acetylcholine response 20 minutes after stimulation onset [14]. Studies using acetylcholine bioassays of cortical samples *in vivo* showed similar results with increased acetylcholine in response to DBS at a 40-minute time interval [15]. Others have shown an increase in acetylcholine release with increasing stimulus intensity and frequency [16,17]. Slice work using whole cell patch of neurons in the entopeduncular nucleus [18] showed that high-frequency stimulation (>100 Hz) of the basal forebrain induced afterdepolarizations, and pharmacological antagonists of acetylcholine muscarinic receptors blocked this response. However, prior work has not explored the pharmacokinetics of acetylcholine in the cerebral cortex in response to deep brain stimulation of the cortical projection nucleus with a time resolution of seconds.

Various methods have been employed to measure acetylcholine in the brain, each differing in sensitivity, accuracy, and temporal resolution. Microdialysis, for example, allows measurement of acetylcholine but requires acetylcholinesterase inhibitors [17,19] and has long sampling times, which may not reflect physiological changes accurately. Despite this, microdialysis has provided important insights into cholinergic dynamics. Amperometric measurement offers faster detection but uses choline as a proxy for acetylcholine [20] and interactions with oxygen can limit interpretation of results [21]. Fluorescence microscopy using the acetylcholine-specific GRAB_ACh3.0_ sensor, a muscarinic receptor coupled to GFP, provides a more direct measurement of acetylcholine [22,23]. It can be targeted to neurons through viral injection with an hSyn promoter, and its extracellular binding domain provides a readout of both synaptic and extra-synaptic acetylcholine concentration. When combined with imaging, this sensor allows for more accurate recording of acetylcholine changes. Fiber photometry has been used to image acetylcholine levels in freely moving animals [23–27]. This method has certain advantages, including rapid acquisition and the ability to monitor acetylcholine in awake moving animals. However, compared to two-photon imaging it has lower spatial resolution, is more susceptible to photobleaching and phototoxicity, and requires invasive probe insertion for deep imaging [24].

To advance our understanding of how the basal forebrain responds to different pulse patterns, we created a preparation that combines deep brain stimulation of the basal forebrain with fluorescence imaging of the GRAB_ACh3.0_ sensor [23] using 2-photon microscopy. In supposing the basal forebrain to be a black box in between the stimulation and the release of acetylcholine in the cerebral cortex, we constructed two experiments. The first applied different frequencies of pulse trains with the same number of pulses to assess frequency selectivity of the basal forebrain. The second applied different lengths of pulse trains at one fixed frequency to assess effects including sustainability of response, saturation of response, and depletion of response. Because we are triggering the release of acetylcholine, we also performed an experiment that combined stimulation with administration of donepezil, an acetylcholinesterase inhibitor used in the treatment of Alzheimer’s disease [28,29]. In theory, acetylcholine levels only increase with synaptic release, and only decrease with its cholinesterase catalyzed hydrolysis. We predicted donepezil would slow hydrolysis, which would result in slower returns to baseline after stimulation.

## Methods

### Test subjects

Male and Female C57/BL6J mice from Jackson Laboratories (Bar Harbor, ME) aged 8-10 months were used. Mice were initially housed in groups of 5 per cage, housed individually after surgery, in a temperature and humidity controlled room on a 12-hour light and dark cycle (06:00-18:00). Animals were given *ad lib* access to food (Teklad Rodent Diet 8604 pellets, Harlan, Madison, WI) and water. All testing was performed during the light cycle portion of their day. All procedures in this study were approved by the Institutional Animal Care and Use Committee of Augusta University. 14 animals in total were used in 35 different experiments for this study.

### Electrode Fabrication

Stimulation electrodes were custom-made based on previously published specifications [12,30]. A Pt/Ir wire with PFA insulation (A-M Systems, Seattle, WA) was stripped of 0.75 mm insulation at the stimulation end and was placed inside a hypodermic tube. The tubing was placed inside polyimide tubing to prevent tissue contact with the stainless steel hypodermic.

### Surgical Procedures

*Viral Transduction* Surgery was performed in a sterile field under 1.5% isoflurane anesthesia. The mouse was placed into a bell jar with 0.5mL isoflurane until loss of righting reflex. The mouse was transferred to a rodent stereotaxic instrument (Stoelting ®, Wood Dale, IL) with a heating pad. Artificial tears ophthalmic ointment (Akorn, Lake Forest, IL) was applied to both eyes. The hair on the mouse scalp was removed with hair remover (Veet). The hairless scalp was swabbed with a povidone-iodine (Dynarex ® Orangeburg, NY) followed by 70% isopropyl alcohol. A 3 mm incision was made to expose the skull. A 0.5 mm craniotomy was made (−2.0mm AP, 2.0 mm ML from bregma). A micropipette was inserted over 3 minutes to a depth of 700μm. Mice were injected with 500nL of (AAV9-hsyn-ACh4.3, Vigene) at 100nL/min using a micropump (World Precision Instruments, Sarasota, FL). After a 10-min wait period, the micropipette was slowly withdrawn from the cortex over 3 minutes. The scalp incision was sealed with a suture. After recovery, the viral vector incubated for at least 3-wks.

*Stimulation Electrode and Optical Window Implantation* Surgery was performed in a sterile field under 1-2% isoflurane anesthesia. The mouse was placed into a bell jar with 0.5mL isoflurane until loss of righting reflex. The mouse was transferred to a rodent stereotaxic instrument (Stoelting ®, Wood Dale, IL) fitted with an anesthesia mask and heating pad set to 37°C. The scalp was disinfected as previously mentioned. Artificial tears ophthalmic ointment (Akorn, Lake Forest, IL) was applied to both eyes. Surgical scissors removed the scalp, and a scalpel blade scraped clean the periosteum. The skull was scrubbed with hydrogen peroxide to remove connective tissue. To form a 3 mm round cranial window, a craniotomy was made using a ¼ round carbide dental bur (Midwest). The brain was irrigated with a sterile saline buffer, and excess bleeding was managed with Vetspon (Elanco). A 4mm diameter #1 glass coverslip which was previously glued to a 3mm diameter #1 glass coverslip (Electron Microscopy Sciences) was placed over the window such that the 3mm inferior section was flush in the craniotomy. Cyanoacrylate was used to seal the coverslip to the skull. Stereotaxic measurements were made in reference to the bregma landmark (3.0mm AP, 1.7mm ML) to target sites for electrode implantation. A 0.5 mm hole was drilled at target sites using a carbide bur (HM1-005-FG, Meisinger, Centennial, CO) and dental drill (Volvere GX, Brasseler, Savannah, GA) until the cranium was breached. An electrode was loaded into a guide tube positioned above the target site at a 22^°^ angle to the horizontal plane and slowly pushed into the cerebrum over 1 min until the back of the electrode became flush with the cortical surface to ensure achievement of 6mm depth (Illustrated in Figure 1B). The angled electrode implantation was used to stimulate the sub-pallidal basal forebrain without obstructing the optical window. Cyanoacrylate was placed at the target site to stabilize the electrode and seal off exposed tissue. The procedure was completed on the right side of the skull ipsilateral to the cranial window. With the same burr, holes were made laterally and anterior on the left for bone screws (Antrin, Fallbrook, CA). Before implanting the anterior bone screw in the left frontal bone, a platinum-iridium wire with 3mm of insulation stripped was inserted to serve as the ground wire. A layer of cyanoacrylate was applied to all screws and electrode sites and left until dry. Methyl methacrylate powder (A-M Systems, Everett, WA) was mixed with its solvent and a thin layer of the mix was applied to the top of the skull with periosteal elevators. A custom made head holder was embedded in the dental cement on the left side of the headcap so that the mouse could be held in place under the microscope. Postprocedural lesions were created in all animals to confirm electrode tip location (Figure 1A). Lesions were created with a 30 μA DC current, electrode negative, for 10 seconds which generated a clearly visible mark on perfused and unstained sections.

**Figure 1.**
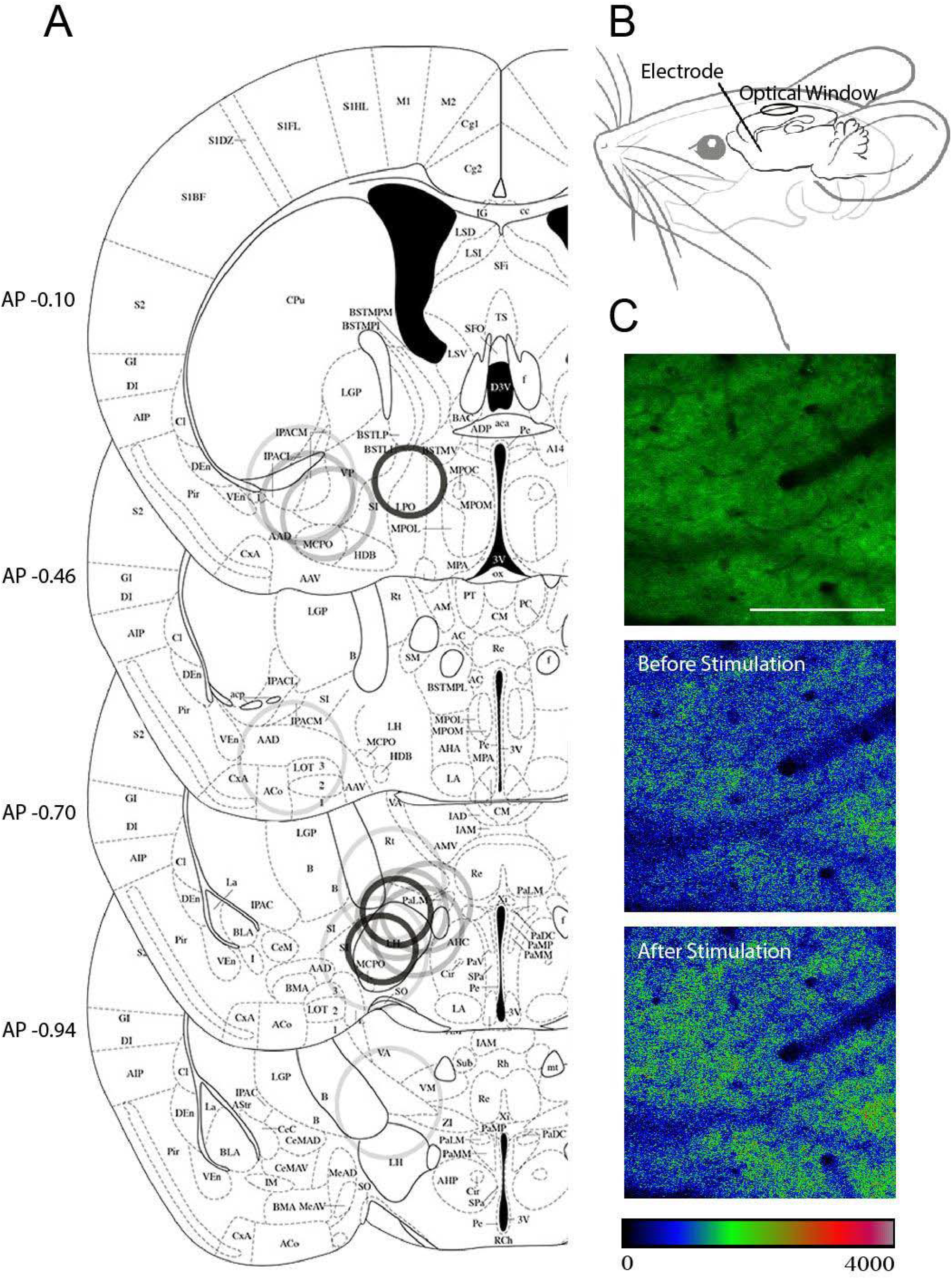
Experimental Procedures. A) Location of electrode tip in animals used in this study showing activating function radius based on current needed to elicit a ∼10% change in fluorescence [34]. Approximate radius of activation is illustrated at electrode tip location with opacity adjusted such that smaller currents are more opaque (≤100uA 95% opacity, 100-125uA 75% opacity, ≥200uA 25% opacity) B) Schematic of surgical preparation from a parasagittal viewpoint C) Example GRAB_ACH 3.0_ GFP Z-projected fluorescence image (Top), single frame color-coded fluorescence intensity before stimulation (Middle) and at the peak after stimulation (Bottom). Scale bar is 100μm. Color gradient scale shows fluorescence intensity (arbitrary units) for Middle and Bottom panels.

### Two Photon Imaging

Fluorescence imaging data was collected using the Ultima 2P-Plus two-photon microscope system (Bruker Corporation) coupled with the Insight X3 laser (Spectra-Physics). Laser power levels were controlled using Pockels Cells (350-80, Conoptics). Imaging was performed at 920 nm with a Nikon 16X (numerical aperture 0.8) water immersion objective (CF175 LWD 16X W), using resonant scanning mode (30 Hz/frame) with 2-frame averaging. Images were collected at 512×512 pixels, typically ∼560×560µm. Figure 1C shows a typical maximum intensity projection (MIP) of a small imaged volume followed by pseudo-colored images before stimulation (middle) and at the peak after stimulation (bottom) indicating differences in fluorescence intensity. Initial experiments indicated that some tissue movement occurred during stimulation. We estimated this motion could be algorithmically corrected (See Data Analysis below) within a 7.5µm depth of tissue. An Electro-Tunable Lens (ETL) was used for rapid Z-plane acquisition. Typically, 6 depth planes separated by ∼1.5 µm were acquired in rapid succession, with each depth plane imaged every 582 ms. Given the axial resolution of our setup (∼1.5 µm), our acquisition parameters seem adequate to ensure no data is lost with potential Z-shifts while avoiding photobleaching that could result from oversampling. Figure 2A shows single images taken at each plane from an example mouse. Z-stacks were collected between 40-50 µm deep from the cortical surface, except in the depth comparison experiment where Z-stacks were collected at 100µm deep. All experiments included infraorbital injection of 40µL Texas Red Dextran (70 kD, 5% in physiological saline, Invitrogen™ D1830) for visualizing vessels and tracking Z-plane shifts.

**Figure 2.**
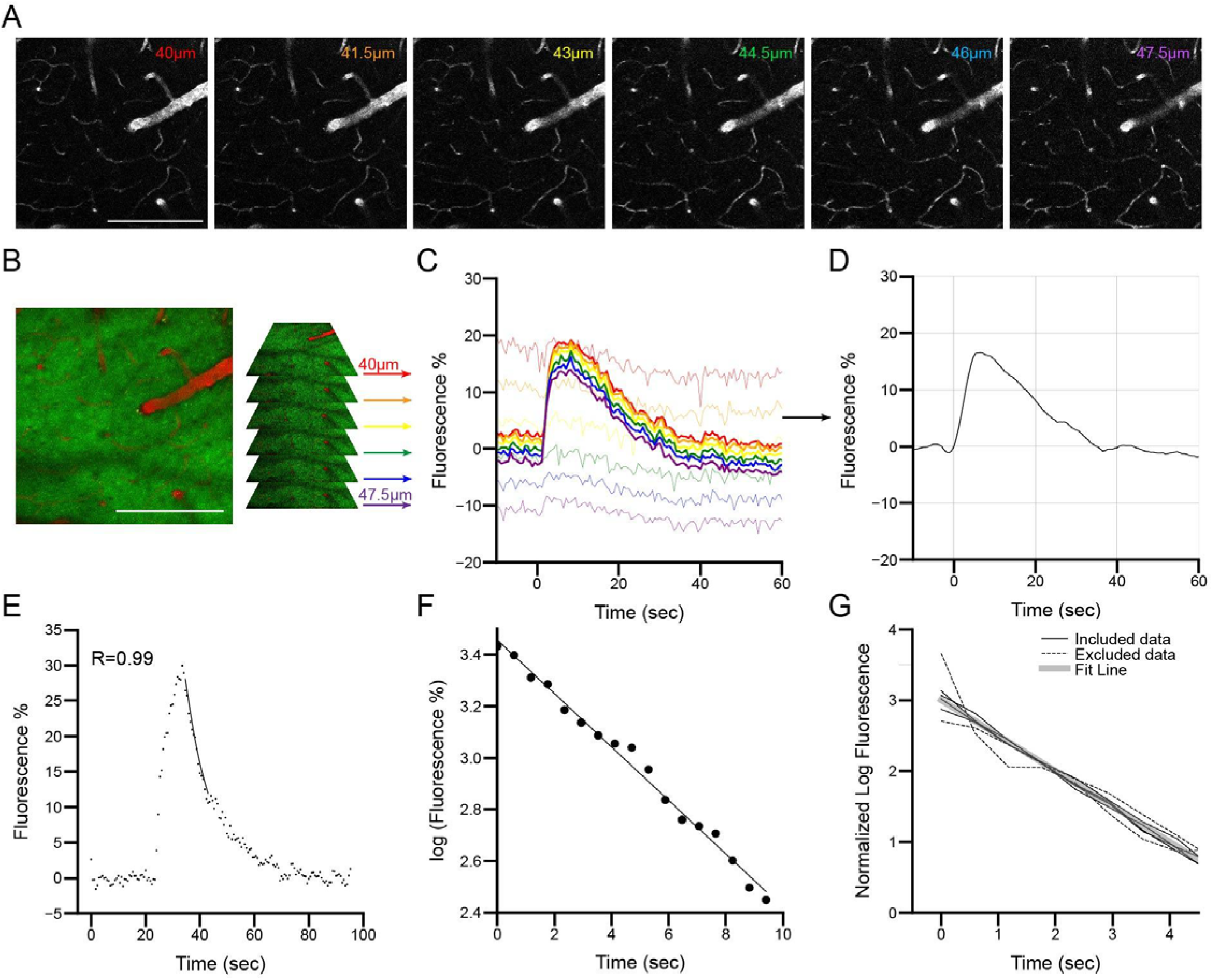
Data Processing. A) Single plane images from a Z-stack labeled with corresponding approximate depth B) Maximum intensity projection (left) and individual depth planes (right) of the imaged volume from A showing blood vessels labeled with Texas Red and neural tissue labeled with GRAB_ACH 3.0_ GFP (green) C) Time course of the average fluorescence change at each depth plane, color coded as in B. Thick colored lines are the green channel planes and thin colored lines are the red channel. D) Green channel trace averaged across corrected Z-planes and smoothed with a Savitzky-Golay smoothing filter. E) Example data without smoothing with an exponential fit overlaid. The data points include time before and after the analyzed recovery period. F) Data points from E mark full frame fluorescence at each time point, log transformed. The line is a best fit.G) Data from individual animals in each experiment and five conditions are log transformed, and then slopes and intercepts adjusted to overlay them on a single line to highlight deviations from an exponential fit. Solid lines show data included in recovery time constant analysis, and dashed lines show data excluded because the exponential fit was inadequate for our criteria. Scale bar is 100μm.

At the beginning of every experiment session, the clarity of the cranial window was evaluated by eye using a 4x objective. Animals with windows that were not clear were terminated from the study.

### Electrode Impedance Measurement

The individual impedances of each electrode were measured to verify successful function and to normalize current used at the electrode tip under voltage control. Animals were used in experiments once per week at the most to give time for the system to recover completely and impedances were measured once per week. Impedance was measured using an oscilloscope (TDS 2012C Tektronix, Beaverton, OR) and custom programmed software that triggered a stimulus isolator (A385 World Precision Instruments, Sarasota, FL) to produce a known 100 μA current square wave. After confirming calibration of the method using a known impedance of a circuit resistor, the same method was applied to the electrode of each mouse head cap. Electrode impedances averaged 10-12 kOhm.

### Brain Electrical Stimulation

Animals received stimulation of the sub pallidal basal forebrain under isoflurane anesthesia while imaging through cranial windows. A 10 μF capacitor was placed in line with the electrode during stimulation to prevent net charge transfer. 600 biphasic unipolar pulses were delivered at different frequencies [20Hz, 60Hz, 100Hz, 130Hz], and 60Hz stimulation was delivered with different pulse train lengths [300 pulses or 5s, 600 pulses or 10s, and 1200 pulses or 20s] in pseudo randomized order. All stimuli were delivered with 0.10ms pulse width. Stimulus pulses were triggered at exactly 240 frames, or 40 Z-stacks into acquisition (23.28 seconds), to ensure that the preparation was stable prior to stimulation. A 300 test pulse train at 60Hz (5 seconds of stimulation) was used at the beginning of each experiment session to determine a starting voltage that would elicit a 5-15% change in fluorescence. Upon success, this voltage was then used in that animal for that day’s experimental session. Pilot studies showed that repeated consecutive stimulation trials had an exhaustive effect on the system, but near full recovery was achieved with a 20-minute wait. Successive stimulation trials were therefore separated by 20 minutes. The activating function for each stimulation pulse was estimated in microns as 313 * sqrt (current/100 microamps) [31,32]. Figure 1A displays estimated activating function based on current used and electrode location to elicit ∼5% change in fluorescence. Reported measures for a given experimental session was performed in the same animal which allows for pairwise comparisons.

### Donepezil administration

Donepezil HCL (Memory pharmaceutical Corporation, Montvale, NJ) was prepared from powder with normal saline (0.85% NaCL, Spectrum Chemical MFG Corp, Gardena, CA) into a 1mg/mL solution and was injected intraperitoneally to animals at either 2mg/kg or 4mg/kg. Animals did not receive more than one dose of donepezil per week nor were they subsequently tested on non-donepezil experiments in the same week. Baseline fluorescence changes to stimulation was recorded for 10s and 60s followed by donepezil administration and a 30 minute [33] incubation period. Then fluorescence was recorded again to stimulation for 10s and 60s.

### Data Analysis

Whole images were analyzed using a custom program designed to track tissue movement. For X,Y, and Z-shifts, a 300×300 pixel frame was selected within the 512×512 red-channel image. This sub-frame was matched by peak cross correlation against all possible X and Y positions within each Z-stack image to find the shift resulting in the best fit, which was subsequently used for correction in the green channel. Figure 2B-D shows the data transformation from images to traces. Figure 2B shows a MIP of both red and green channels followed by color coded z-planes that correspond to the plotted time course in Figure 2C for both the red (thin lines) and green channels (thick lines), and finally the algorithmically corrected trace in Figure 2D. The quality of each run was scored from 0 - 5 plane shifts where 0 denotes no shifts on the Z-plane and 5 denotes shifts of 5 planes. On average, runs scored 0.8 on Z-shifting with a mode of 0.5 indicating a generally stable preparation and no runs were excluded on this basis. When they occurred, shifts in the Z axis peaked 10 seconds after stimulation onset, and were never more than 1 Z-plane more than 30 seconds later. Shifts were highest in the period 5 to 20 seconds after stimulation onset.

Acetylcholine based fluorescent “green” data was normalized to the average value over the 10s preceding the onset of stimulation and is presented as %-change from this baseline. All data in this work are whole frame Z-Stack averages of the green channel taken from the tracked tissue, and all Z-planes that remained in the imaging zone were averaged together for each trial.

Once Z-shift corrected data was acquired from the images the following were calculated for each run:

- *Area Under the Curve (AUC)* Area under the curve of the fluorescence trace was calculated as a metric of acetylcholine release during stimulation. We integrated the AUC between the onset of stimulation and 10s after stimulation offset for all experiments. For Donepezil experiments and the 20Hz condition where some traces returned to baseline in less than 10s following stimulation offset, AUC was integrated only over the stimulation duration.
- *Peak Fluorescence and Time to Peak* Peak fluorescence was calculated by averaging every 3 data points and finding the highest value. Time to Peak is the time taken to achieve this peak from stimulus onset.
- *Time Constant* Time constants were calculated to estimate recovery time. In principle, after the response peaked, the response, if it were time invariant, would exponentially decay back to baseline. A fit to an exponential with a high explained variance, or high *R*^*2*^, would be consistent with the hydrolysis of acetylcholine being primarily determined by its concentration. The analyzed data were in an 8-10 second period after peak was reached, which occasionally was as much as three seconds after stimulation ended. The asymptotic baseline at the end of the run was subtracted from the data. The data were log transformed (Figure 2F) and fit to a line. The slope of that line was the negative inverse of the time constant, or the time over which the response would decay from peak to 1/e of its peak value. The correlation coefficient was calculated, in log space, between the best fit line and the data, and a correlation *R* greater than 0.75 was used for quality control (Figure 2E). 85.3% (105/123) of the data were included for Time Constant analysis (Figure 2G).

### Statistical Analysis

ANOVAs and other statistical metrics in this work were calculated using the Python library Pingouin. When paired data were used, post hoc tests were run on the computed differences between groups and compared to 0. P-values were adjusted with Bonferroni correction. When data were not paired, Tukey’s post-hoc was used. Two tailed Student’s t-tests were used when comparing only two groups.

## Results

### Effect of Frequency on Acetylcholine response

The first experiment assessed frequency selectivity of the basal forebrain response to pulse train stimulation by applying the same number of pulses at different repetition rates. Eleven animals were used, and each stimulus contained 600 pulses. Tested frequencies were 20Hz, 60Hz, 100Hz, and 130Hz. In each experiment, the same imaging position was used for each frequency. Twenty-minute recovery periods were also used between runs. An example from one animal showing the response to different frequencies is displayed in Figure 3A. Example traces are smoothed with a Savitzky-Golay smoothing filter. In this figure, and in all data in this work, full frame fluorescence from single trials was used to quantify the acetylcholine response (∼1.7Hz sampling rate) in the tracked tissue volume (see Methods).

**Figure 3.**
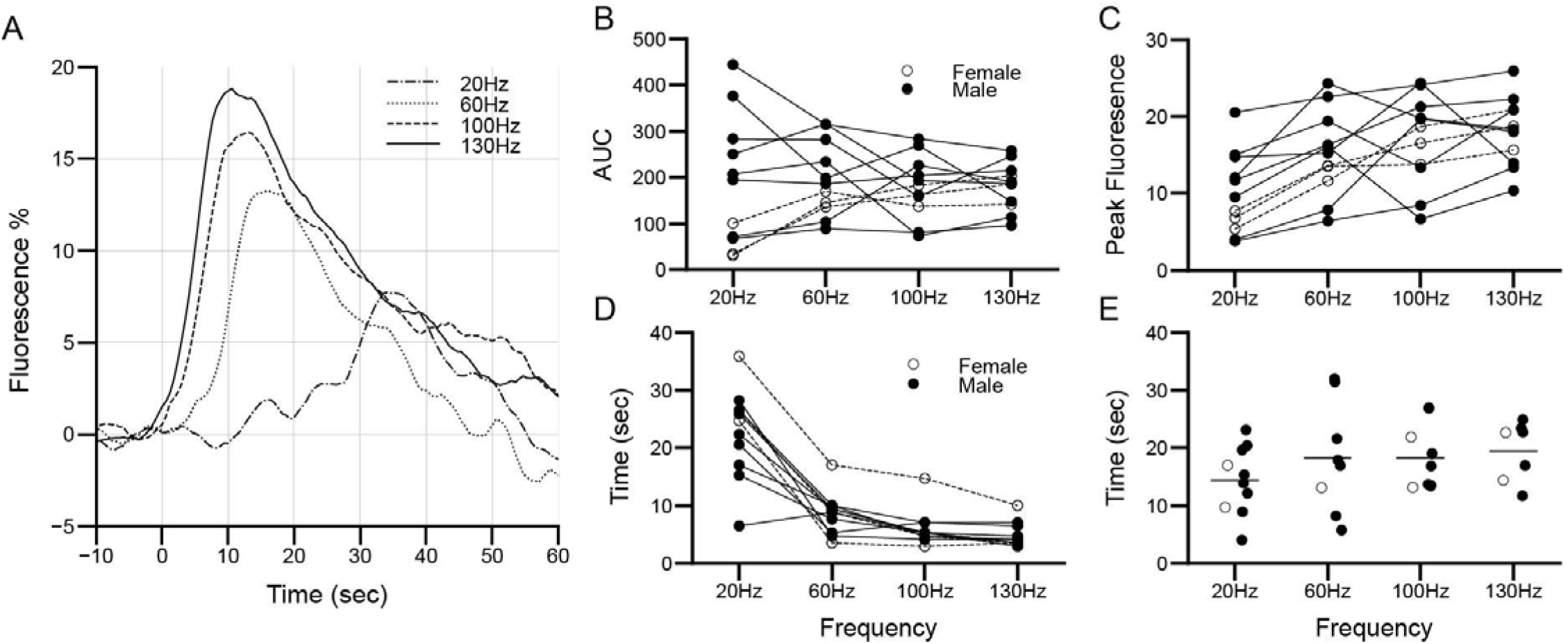
Effect of frequency on Acetylcholine response. A) Example traces from a frequency experiment. Lines show whole frame mean fluorescence changes from baseline in response to stimulation pulse trains beginning at time 0 B) Area Under the Curve. Summed fluorescence increases over baseline in response to each frequency. C) Peak fluorescence amplitude D) Time to peak (n=11) E) Time constant (n=8-11). Each line and dot represent an individual animal. (Female n=3, Male n=8).

To estimate the total cholinergic response, we calculated the area under the curve (AUC) from stimulus onset until 10 seconds after stimulus offset (see Methods). An ANOVA comparing the AUC from different frequencies of stimulation did not find a significant effect [Figure 3B: F(3, 40) = 0.094, p = 0.962].

Higher frequencies of stimulation resulted in significantly higher peak acetylcholine induced fluorescence compared to the 20 Hz condition (Figure 3C). An ANOVA on the peak fluorescent response found a significant impact of frequency. [F(3, 40) = 4.803, p = 0.006]. Post-hoc tests used a paired statistical approach and used a Bonferroni corrected p-value of 0.008. The 20Hz peaks were lower than those at 60Hz [t(10) = 5.21, p = 0.0003], 100Hz [t(10) = 4.00, p = 0.003], and 130Hz [t(10) = 5.21, p = 0.0004], with no other differences.

The Time to Peak was faster for higher frequencies than it was for the 60 Hz and 20 Hz conditions [Figure 3D: F(3, 40) = 34.16, p < 10^−11^]. Post hoc, paired, tests showed 20Hz was slower than all other frequencies tested: 60Hz [t(10) = 6.29, p = 9e^-5^], 100Hz [t(10) = 7.74, p = 2e^-5^], and 130Hz [t(10) = 8.28, p = 9e^-6^], and that 60Hz was slower than 100Hz [t(10) = 4.13, p = 0.002] and 130Hz [t(10) = 4.45, p = 0.001] with no other differences.

Time constants of recovery were not significantly impacted by frequency (Figure 3E), and the average across all frequencies was 17.47 seconds. To determine the recovery of the response, we calculated the time constant after stimulation ceased. If the rate of clearance of acetylcholine is dependent only on its concentration the recovery function would be purely exponential. Therefore, an 8 or 10 second segment after the stimulation response peaked was fit to an exponential decay (see Methods). No significant differences were found between groups [F(3, 30) = 0.983, p = 0.414]. Data were plotted as independent groups instead of paired lines because recoveries that were not well fit by an exponential were not included, which means that data for this analysis was not strictly paired as it was for the other analyses.

To determine if there was any effect of sex, with consideration that the group sizes are smaller for these analyses, we subdivided the data into male and female groups. No significant effects were found when comparing any of the parameters with t-tests.

The remaining experiments were all performed at 60Hz.

### Effect of Number of Pulses on Acetylcholine response

The second experiment assessed the impact of pulse train lengths on basal forebrain cholinergic response in cortex. All experiments were conducted with 60Hz pulse trains so longer pulse train lengths also correspond to more pulses. Nine animals comprised the data set examining the effects of 300 (5 seconds), 600 (10 seconds), and 1200 (20 seconds) pulse train lengths. Again, the entire set of stimulus durations were collected from the same imaging location, and data for most analyses were paired. An example from one animal showing the response to different pulse train lengths are shown in Figure 4A. Again, each line in Figure 4A is defined in a single trial with samples taken in the tracked tissue volume.

**Figure 4.**
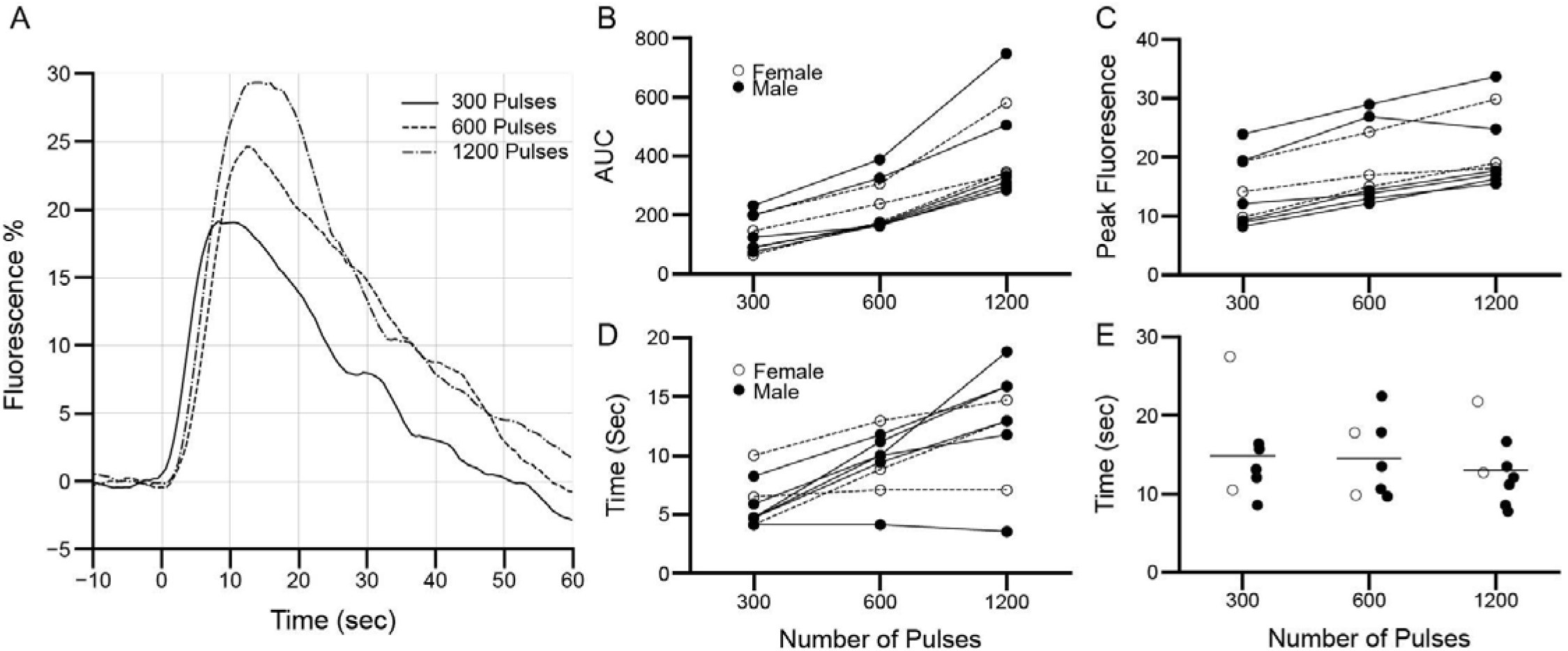
Effect of Number of Pulses on Acetylcholine response. A) Example traces in response to 60 Hz pulse trains of different lengths. B) Area Under the Curve C) Peak fluorescence amplitude D) Time to peak (n=9) E) Time constant (n=7-8). Each line and dot represent an individual animal for panels B-D. (Female n=3, Male n=6).

Total cholinergic impact of stimulation increased with longer pulse train lengths. An ANOVA comparing areas under the curve with different stimulation times showed a significant effect. [Figure 4B: F(2, 24) = 14.864, p = 6e^-5^]. As in previous experiments, post-hoc tests used a paired statistical approach and used a Bonferroni corrected p-value of 0.017. Differences between all groups were significant [1200 vs 600 pulses: (t(8) = 6.64, p = 0.0002), 1200 vs 300 pulses: (t(8) = 7.74, p = 6e^-5^), 600 vs 300 pulses: (t(8) = 8.81, p = 2e^-5^)]. Mean AUC were 300 pulses: 134.93, 600 pulses: 232.56, 1200 pulses: 415.64, therefore, 600 pulses AUC was 1.7 times larger and 1200 pulses AUC was 3.1 times larger than 300 pulses. All cross-condition comparisons indicated sublinear summation i.e., the 600 pulse length response AUC was less than twice the 300 pulse length response, and the 1200 pulse length response less than double the 600 pulse length response.

Longer pulse train lengths impacted the amplitude of acetylcholine responses, as shown in Figure 4C. An ANOVA on peak fluorescence amplitude showed a near significant effect of pulse train length [Figure 4C: [F(2, 24) = 3.24, p = 0.057]. As the data within each experiment appeared strongly dependent on duration, while the data across experiments were more variable, we performed another ANOVA on this data after normalizing each within-experiment triplet to a mean of 1 and found significant impacts of pulse train length on peak amplitude [F(2,24) = 93.919, p = 4e^-12^] with post hoc tests showing significant differences for all comparisons [1200 vs 600 pulses: (t(8) = 5.44, p = 4e^-5^), 1200 vs 300 pulses: (t(8) = 13.61, p < 10e^-6^), 600 vs 300 pulses: (t(8) = 8.18, p = 6e^-8^)]. The longer pulse train lengths resulted in high peak responses. Mean Peak responses were 300 pulses: 113.86, 600 pulses: 118.35, 1200 pulses: 121.31, therefore, the 600 pulse Peak was 1.04 times larger and 1200 pulse Peak was 1.07 times larger than 300 pulses. Again, all cross-condition comparisons indicated sublinear summation.

Time taken to reach peak responses were longer with longer pulse train lengths. An ANOVA on Time to Peak found a significant effect of pulse train length [Figure 4D: F(2, 24) = 9.162, p = 0.001]. Post hoc tests again showed significant differences between groups [1200 vs 600 pulses: (t(8) = 3.32, p = 0.011), 1200 vs 300 pulses: (t(8) = 4.50, p = 0.002), 600 vs 300 pulses: (t(8) = 5.06, p = 0.001)]. All peak times in the 1200 pulse train (20 second) group were earlier than 20 seconds. Mean Time to Peak were 300 pulses: 5.88s, 600 pulses: 9.47, 1200 pulses: 12.61.

Time Constants were not affected by duration of stimulation, and the average across all pulse train lengths was 14.05 seconds. An ANOVA on Time Constant showed no significant effect [Figure 4E: F(2, 19) = 0.261, p = 0.773].

Again, we subdivided the data into male and female groups and no significant effects were found.

### Effect of Donepezil on Acetylcholine response

Donepezil is an acetylcholinesterase inhibitor that we expected would slow recovery times because it inhibits acetylcholine hydrolysis. We examined the effect of 2mg/kg and 4mg/kg Donepezil on acetylcholine release to basal forebrain stimulation in eleven animals in total. Five animals were tested in the 2mg/kg group and six animals were tested in the 4mg/kg group. We first imaged baseline acetylcholine responses to 10s at 60Hz. We then injected donepezil (i.p.) and allowed 30 minutes for incubation before we imaged again at the same location with the same stimulation protocol. The data for most analyses were paired and representative example data are shown in Figure 5A&E.

**Figure 5.**
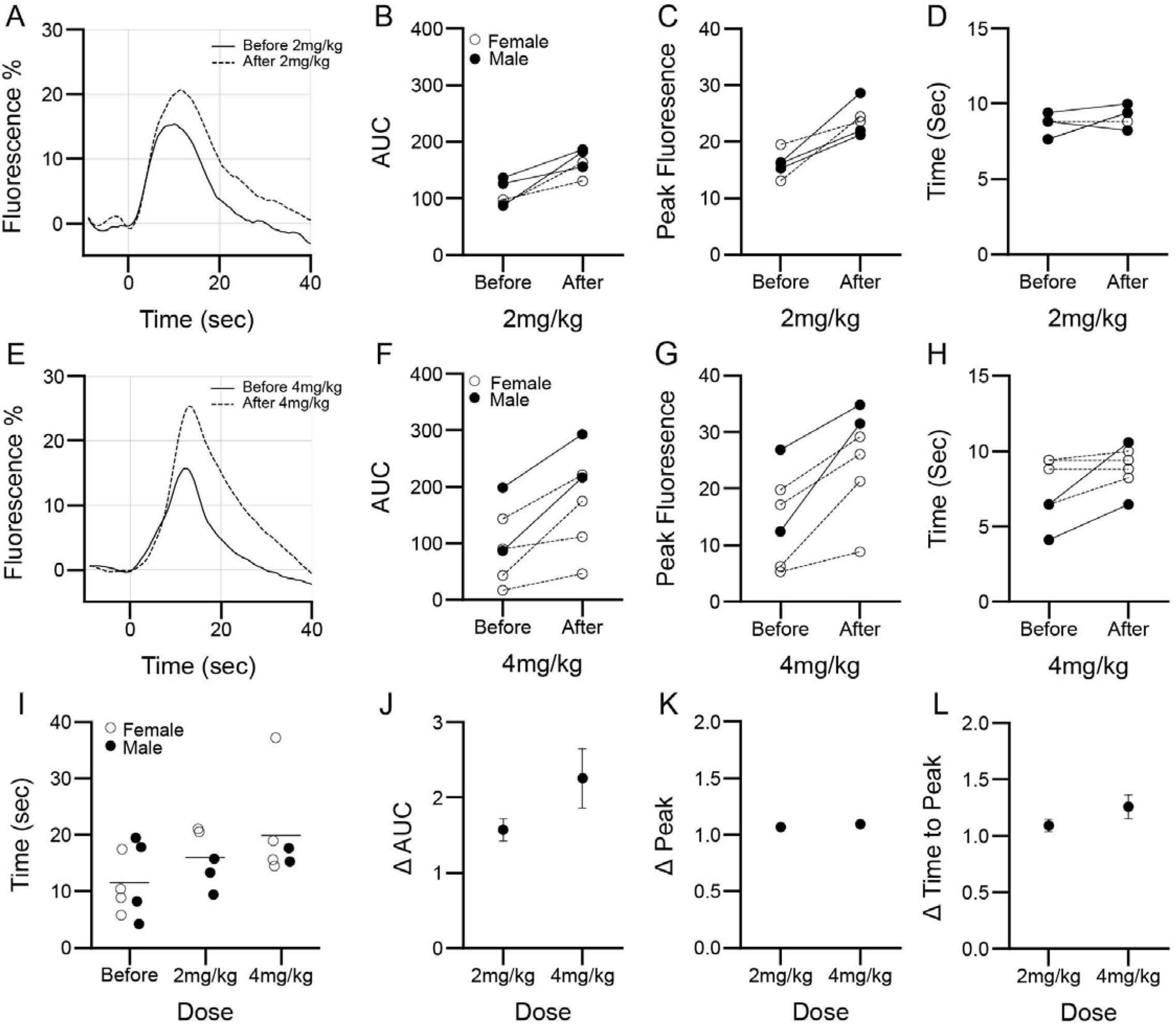
10s stimulation with Donepezil. A-D) Responses to 2mg/kg Donepezil (n=5, Female n=2, Male n=3) A) Example trace B) Area under the curve C) Peak Fluorescence amplitude D) Time to peak. The values 7.644 before and 9.408 after are actually two points, one male and one female. E-H) Responses to 4mg/kg (n=6, Female n=4, Male n=2) E) Example trace F) Area under the curve G) Peak Fluorescence amplitude H) Time to peak I-L) 2 and 4mg/kg (n=5-11) I) Time constant J) Ratio change in Area under the curve K) Ratio change in Peak Fluorescence L) Ratio change in Time to Peak. Each line and dot represent an individual animal.

Donepezil administration increased integrated acetylcholine response with both doses. A t-test on AUC between before and after administration of each dose showed significant effects [Figure 5B: 2mg/kg: t(4) = 4.499, p = 0.011, Figure 5F: 4mg/kg: t(5) = 4.143, p = 0.009]. We normalized the paired data to their individual baseline and compared both doses. Notably, although there was no statistical difference [Figure 5J: t(9) = 1.36, p = 0.208], AUC for 2mg/kg had a mean of 1.573+/- 0.1485 and 4mg/kg had a mean of 2.2563 +/- 0.3969, which constitute 57.3% and 125.6% increases respectively.

Peak amplitude acetylcholine responses increased with donepezil administration with no effect on Time to Peak. Peak responses were significantly higher for both doses [Figure 5C: 2mg/kg: t(4) = 4.800, p = 0.009, Figure 5G: 4mg/kg: t(5) = 4.715, p = 0.005]. When normalized, there was no difference between doses [Figure 5K: t(9) = 0.952, p = 0.366] with mean of 1.07+/-0.01 and 1.09+/-0.02 representing 7% and 9% increases for 2mg/kg and 4mg/kg respectively. Time to Peak was not significantly impacted by these doses of Donepezil [Figure 5D: 2mg/kg: t(4) = 1.50, p = 0.208, Figure 5H: 4mg/kg: t(5) = 2.24, p = 0.076] with no difference when normalized [Figure 5L: t(9) = 1.18, p = 0.269].

Time Constants were not affected by either dose of donepezil, with an average of 15.53 seconds. A Dose x Time Constant ANOVA was not significant [Figure 5I: F(2,16) = 2.70, p = 0.0979].

There was no effect of dose across any of the parameters extracted [AUC: t(9) = 1.36, p = 0.208, Peak: t(9) = 0.952, p = 0.366, Time to Peak: t(9) = 1.78, p = 0.269].

No sex differences were observed except for the Time Constant in the 2mg/kg condition (Figure 5I: p = 0.045).

To determine if pulse duration interacts with the effects of donepezil, we examined the effect of 2mg/kg and 4mg/kg Donepezil on 60s of stimulation at 60Hz (3600 pulses) in the same animals as in Figure 5. Representative data are shown in Figure 6A&E.

**Figure 6.**
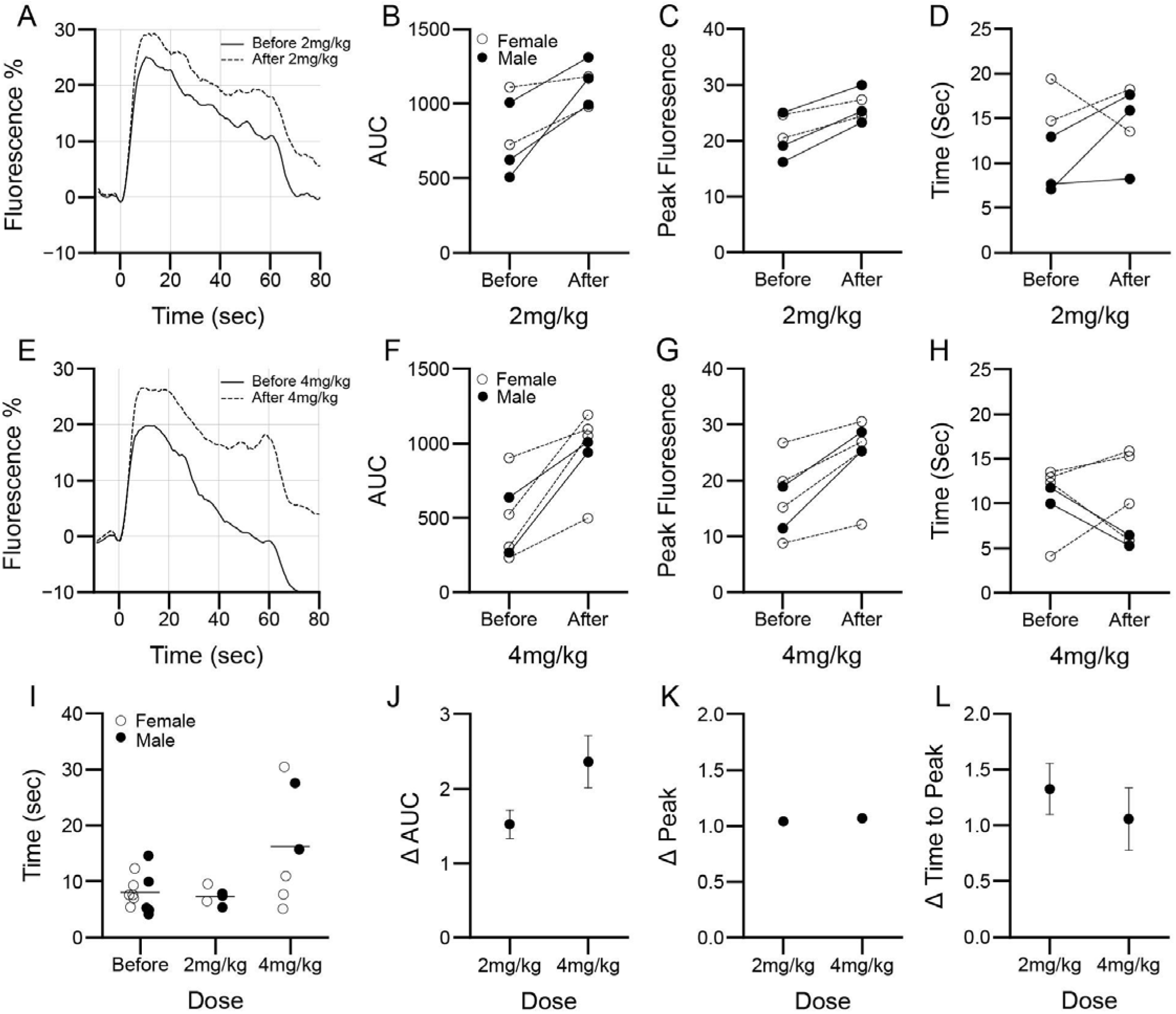
60s stimulation with Donepezil. A-D) Responses to 2mg/kg Donepezil (n=5, Female n=2, Male n=3) A) Example trace B) Area under the curve C) Peak Fluorescence amplitude D) Time to peak. E-H) Responses to 4mg/kg (n=6, Female n=4, Male n=2) E) Example trace F) Area under the curve G) Peak Fluorescence amplitude H) Time to peak I-L) 2 and 4mg/kg (n=5-11) I) Time constant J) Ratio change in Area under the curve K) Ratio change in Peak Fluorescence L) Ratio change in Time to Peak. Each line and dot represent an individual animal.

Donepezil administration again increased integrated acetylcholine response with both doses. A t-test on AUC between before and after administration of each dose showed significant effects [Figure 6B: 2mg/kg: t(4) = 3.457, p = 0.026, Figure 6F: 4mg/kg: t(5) = 4.99, p = 0.004]. Normalized paired data again showed no difference between doses [Figure 6J: t(9) = 1.78, p = 0.108] but we note that 2mg/kg averaged 1.5236 +/- 0.1905 and 4mg/kg averaged 2.361+/-0.3533 which represents a 52% and 136% increase respectively.

Peak acetylcholine responses increased with donepezil administration and again had no effect on Time to Peak. T-tests on Peak responses were significant for both doses [Figure 6C: 2mg/kg: t(4) = 6.369, p = 0.003, Figure 6G: 4mg/kg: t(5) = 4.859, p = 0.005]. When normalized, there was no difference between doses [Figure 6K: t(9) = 1.58, p = 0.149] with mean of 1.04+/-0.01 and 1.07+/-0.01 representing 4% and 7% increases for 2mg/kg and 4mg/kg respectively. Time to Peak analysis did not reach significance [Figure 6D: 2mg/kg: t(4) = 0.962, p = 0.391, Figure 6H: 4mg/kg: t(5) = 0.466, p = 0.066] with no difference when normalized [Figure 6L: t(9) = 0.656, p = 0.528].

Time Constant for 60s stimulation was affected by donepezil administration, with an average of 10.10 seconds. An ANOVA on dose showed a significant effect [Figure 6I: F(2, 19) = 4.424, p = 0.026] with Tukey’s post hoc tests showing only a difference between 4mg/kg and Before [t(16) = 2.729, p = 0.034].

There was no effect of dose across any of the parameters extracted with 60s stimulation as well [AUC: t(9) = 1.78, p = 0.108, Peak: t(9) = 1.58, p = 0.149, Time to Peak: t(9) = 0.656, p = 0.528].

No sex differences were observed.

### Acetylcholine response to 10s stimulation at 40µm and 100µm depth

To keep the laser power relatively low (35-45mW) during imaging we collected most of our data at a depth between 40 and 50 μm. To confirm that the dynamics were likely the same at lower depths, in four animals we ran an additional experiment collecting images at 100µm with the laser power turned up (55-70mW) to achieve similar baseline fluorescence as at 40µm. Representative traces are shown in Figure 7A.

**Figure 7.**
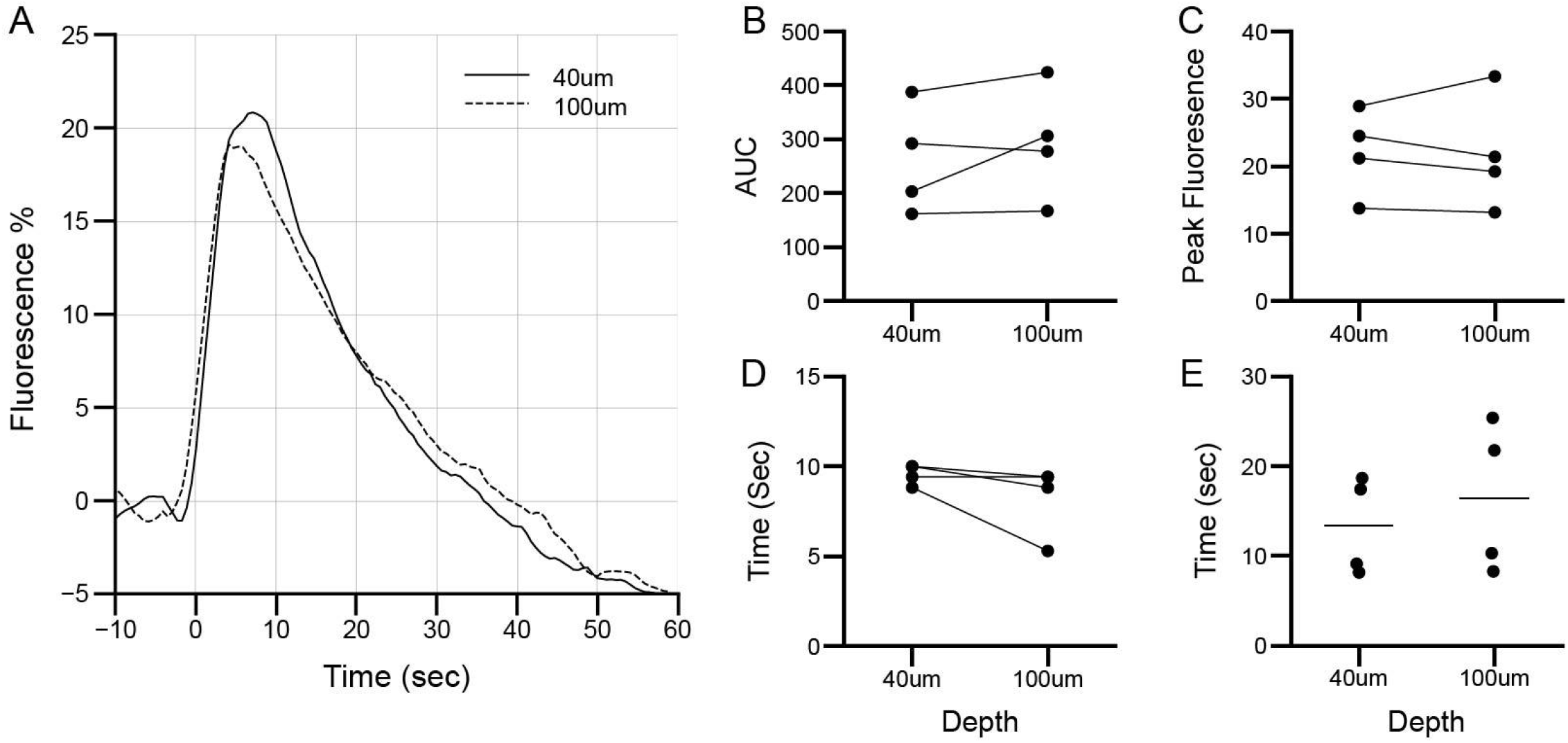
Depth comparison. A) Example traces 40µm and 100µm. B) Area Under the Curve C) Peak Fluorescence D) Time to peak E) Time constant. (n=4) Each line and dot represents an individual animal.

Results were the same regardless of depth chosen. We found no differences in AUC [Figure 7B: t(3) = 1.267, p = 0.294], Peak Fluorescence [Figure 7C: t(3) = 0.189, p = 0.862], Time to Peak [Figure 7D: t(3) = 1.711, p = 0.186], nor Time Constant [Figure 7E: t(3) = 1.779, p = 0.173] which had an average of 14.88 seconds across both depths.

## Discussion

There is increasing interest in the use of DBS of the nucleus basalis of Meynert as a therapeutic modality in Alzheimer’s disease, Parkinson’s disease, and other forms of dementia. This interest is primarily based on considerable work in animal models indicating that DBS increases the release of acetylcholine in the cortex, enhances cerebral blood flow, induces release of several neuroprotective factors, facilitates plasticity of cortical and subcortical receptive fields, and improves the performance of memory-related behavioral tasks (reviewed, [35]). To date a small number of DBS clinical trials have been conducted with some positive, albeit inconclusive, and in some cases equivocal findings. In both the animal models and human studies, a variety of stimulation parameters have been utilized (reviewed, [36]) and it is clear that such parameters have not been optimized thus far. The purpose of the study described here was to employ a novel method for accurately assessing cortical acetylcholine release to determine optimal stimulation parameters as well as evaluate the effect of donepezil, the most commonly prescribed pro-cognitive agent for dementia, on the effect of DBS.

We utilized a wild-type mouse preparation that incorporated deep brain stimulation of the sub pallidal basal forebrain with the GRAB_ACH3.0_ sensor [22,23] for *in vivo* imaging of acetylcholine dependent signal changes. The sensor is virally packaged with an hSyn promoter to specify neuron expression and was injected in somatosensory cortex. The sensor receptor is expressed on the cell membrane and therefore changes in fluorescence indicate extrasynaptic and synaptic acetylcholine levels. This sensor is specific to acetylcholine as opposed to other molecules such as choline [22]. The binding curve for acetylcholine detection with this sensor is sensitive between 0.01-10 μM [23], and the sensor saturates with acetylcholine concentrations near 100 μM with 300% increases in fluorescence. We observed fluorescent changes of 5 to 30%, which suggests acetylcholine levels were in the lower end of the dynamic range for the sensor, likely less than 1 μM.

We examined the cortical acetylcholine response to different pulse patterns using 2-photon imaging and deep brain stimulation. We tested different frequencies (20Hz, 60Hz, 100Hz, and 130Hz) with the same number of pulses, different number of pulses (300, 600, 1200) corresponding to 5, 10, and 20 seconds of stimulation at 60Hz frequency, and the effect of two doses of Donepezil (2mg/kg and 4mg/kg), an acetylcholinesterase inhibitor. Experimental manipulations were performed in the same animal in the same imaging session.

We found integrated acetylcholine responses (AUC) increased with longer stimulation times and donepezil administration but were unaffected by stimulation frequency. Acetylcholine response peak amplitudes increased with higher frequency, longer stimulation times, and donepezil administration. Time taken to achieve these peaks decreased with higher frequencies and increased with longer stimulation times. There was no change in Time to Peak with donepezil administration. These findings were similar between 40 µm and 100 µm cortical depths. Experiments were conducted in male and female mice but this initial work was not focused on sex-specific effects and was underpowered to find them. Additionally, acetylcholine is involved in arousal and wakefulness therefore, the kinetics of acetylcholine responses may be different in the awake animal. These dynamics are also reflective of the somatosensory cortex. Cholinergic projections from the basal forebrain innervate the entire cortical mantle except the medial temporal regions [37]. Some innervated regions are not feasible to image with the current preparation but are being explored by other groups using fiber photometry [23,26,27].

The cholinergic response to subpallidal basal forebrain is sensitive to 60Hz pulse train lengths. The responses were not differentiated by frequencies above 60Hz when the number of pulses was constant. Integrated fluorescent responses were therefore only heavily modulated by the number of pulses. Moreover, at 60Hz and above, peak fluorescent responses were not different. Studies using microdialysis showed that at a one second repetition period and 10 pulses, 100Hz was more effective than 10, 50, and 200Hz at eliciting an acetylcholine response over a 20 minute period [17]. In addition, 60Hz stimulation appears to elicit more effective neuronal responses in primate basal ganglia [51]. We find a significant increase in peak acetylcholine response at 60Hz with an increasing number of pulses. Notably, time to achieve peak responses are under 20 seconds at 60Hz and higher frequencies. Time to Peak remained below 20s for all stimulation durations including for 60s (3600 pulses). Conversely, at 20Hz, Time to Peak, on average, was more than two times higher and with a larger variability in the same animals (22.665 +/- 2.338 vs 4.864-8.606 +/-0.645-1.085). The 20 Hz stimulation also produced much lower peak responses than higher frequencies. These findings suggest that 20Hz stimulation of the basal forebrain may not be optimal for eliciting high peak cholinergic responses. Others have found low frequency intermittent stimulation of the basal forebrain to be effective in improving memory deficits [38], indicating that consideration should be given to stimulation patterns for on/off periods.

From these data, we infer that the minimum stimulation to elicit the highest peak acetylcholine response is 600-720 pulses delivered in up to 12 seconds. In the 300 pulse train condition (5 seconds), timing of peak responses averaged 7 seconds, 2 seconds after stimulation ended. In the 1200 pulse train condition (20 seconds) peak timing averaged 12 seconds. From these two observations we infer that 10-12 seconds of stimulation at 60Hz will consistently result in maximal peak acetylcholine levels. To determine instead the highest acetylcholine response averaged over minutes may require some combination of phasic on and off periods during recovery rather than a sustained low rate.

Peak responses were also increased with donepezil administration. This finding suggests that acetylcholinesterase, responsible for breaking down acetylcholine, contributes significantly to maintaining the ceiling which may be important in modulating the modulatory effects of acetylcholine. Modifying peak values for acetylcholine release is of interest considering that acetylcholine signaling through the α7 nicotinic receptor is neurotrophic and metabotropic [39–43]. This receptor has a much higher EC50 compared to α4β2 [44] which is also involved in maintaining arousal states [4] and competes with the α7 receptor for acetylcholine. Higher peak responses would be expected to increasingly activate the lower affinity neurotrophic α7 nicotinic receptor.

Recovery time was not affected by frequency, number of pulses, or low dose donepezil administration. As reductions in acetylcholine levels are thought to be nearly exclusively the product of hydrolysis, clearance from the interstitial space is governed by acetylcholine hydrolysis followed by choline transport. We modeled the clearance as an exponential decay function [45]. Such a function could govern clearance if hydrolysis was primarily determined by concentration of acetylcholine i.e., if the process was time invariant. Evidence for fast-acting synaptic transmission of acetylcholine exists at the neuromuscular junction where acetylcholinesterase is abundant and terminates acetylcholine responses with sub millisecond timing [46]. Long recovery times are speculated to be evidence for volume transmission, or diffusion through the interstitial space and activation of extrasynaptic receptors, in cortex [5,47]. Notably, the average recovery times in our study were 14-18 seconds. The observed acetylcholine levels are likely reflective of interstitial fluid levels rather than synaptic levels due to GRAB_ACh3.0_ localization [23], which further supports a mechanism necessary for volume transmission. Others have found recovery times between 8-10 seconds in prefrontal cortex in response to foot-shock using fiber photometry [48]. Recovery time was hypothesized to be affected by donepezil administration, but this was only observed with a high dose of donepezil (4mg/kg) after 60 seconds of stimulation. Continuous treatment with donepezil might alter recovery times as expected but in acute settings it appears to modulate acetylcholine levels rather than changing recovery time constants. Recovery time may also be affected by age and disease conditions where cholinergic tone is impaired. Microdialysis measurements of acetylcholine responses in freely moving animals in the amygdala showed a return to baseline after an hour [49,50]. On the other hand, amperometric detection, which has a higher time resolution than microdialysis, of *in vivo* cholinergic transmission in anesthetized animals found that exogenous choline was cleared in 10 seconds [20].

The work presented in this paper is a description of first order cholinergic responses in the somatosensory cortex to various stimulation parameters in the basal forebrain. Findings have immediate potential to inform tuning of deep brain stimulation parameters in patients with Alzheimer’s Disease and Parkinson’s disorder. Surprisingly responses were not frequency selective and seemed to deplete in less than 20 seconds.

## Author CRediT

**Khadijah Shanazz:** Investigation, Writing - Original Draft, Formal Analysis, Visualization, Data Curation. **Kun Xie:** Methodology, Supervision, Investigation. **Tucker Oliver:** Methodology, Investigation. **Jamal Bogan:** Methodology. **Fernando Vale:** Methodology. **Jeremy Sword:** Conceptualization, Methodology, Writing - Review & Editing. **Sergei Kirov**: Conceptualization, Methodology, Writing - Review & Editing. **Alvin Terry:** Methodology, Writing - Review & Editing. **Philip O’Herron:** Methodology, Software, Validation, Supervision, Resources, Writing - Review & Editing. **David Blake:** Conceptualization, Methodology, Software, Validation, Resources, Writing - Review & Editing, Supervision, Project administration, Funding acquisition.

## Acknowledgements

All data shown was collected in the laboratory of Philip O’Herron. Preliminary data for this project were collected using the Department of Neuroscience and Regenerative Medicine 2-Photon microscope. The authors would like to thank Kendyl Pennington for support with coding. This work was supported by the National Institutes of Aging/National Institutes of Health [RF1AG060754] and National Institutes of Neurological Disorders and Stroke/National Institutes of Health [R01NS126920] and [R01NS083858].

